# scFair: Geometry-Aware Gene Budgets and Same-Rank Extension for Highly Variable Gene Selection

**DOI:** 10.64898/2026.08.08.743679

**Authors:** Zhao Li, Aaron W. James, Shengxuan Li

## Abstract

**Background:** Highly variable gene (HVG) selection begins almost every single-cell RNA-seq analysis. While ranking formulas have been compared extensively, the integer gene budget at which any ranking must be truncated is typically left to the user and habitually fixed near 2,000. Relying on such a convention carries hidden costs—lists that are too short erase subtle structure, whereas lists that are too long add noise and computational overhead. Moreover, because global rankings measure variance across all cells, markers for rare populations often lose the “variance vote count” to dominant bulk variation, leading to an unfair feature allocation at the hard cutoff. Whether this convention is defensible, and whether the budget and tail can be set from data without disturbing the ranking, has not been examined systematically.

**Results:** Under a frozen seurat_v3 ranking, k-sweeps across 18 labeled datasets show that n = 2,000 is ARI-optimal on 1 of 18 datasets and that the best available budget is worth a mean ARI gain of +0.033 over it, establishing cardinality as a real and largely unexploited design axis. We present scFair, a Scanpy-compatible HVG layer that automates list length alone: geometry-aware auto_n sets a base size k from multi-seed density and stability features of an intermediate embedding (trading a modest, intentional compute increase for a safer data-driven default), and a same-rank append step acts as a conservative safeguard against cutoff unfairness by adding a short near-miss tail. The ranking is never recomputed or reweighted. On the 18-dataset panel, the default path improved Leiden–label agreement over HVG@2000 (median ΔARI = +0.016; 13/5; Wilcoxon P = 0.0077) and outperformed the neighborhood-based selector triku at author defaults on 15/18 datasets (median +0.024; P = 0.004), while triku did not improve on HVG@2000. Controls locate the effect: cell-number-only rules do not beat HVG@2000, an FDR-chosen length imposed on the frozen ranking is flat, and a fixed HVG@2200 default is not a general substitute because it cannot produce the short lists that compact matrices call for.

**Conclusions:** A fixed budget near 2,000 HVGs is frequently suboptimal, and list cardinality is a separable design axis that can be automated without changing the ranking formula. Effect sizes are modest, the short-list branch rests on four datasets, and rule thresholds were developed with partial overlap to the evaluation panel.

## 1. Introduction

Single-cell RNA-seq analysis almost always begins by restricting the feature space to highly variable genes (HVGs). Established recipes rank genes by excess variance or a related statistic after accounting for the mean–variance trend ^1–3^, and mainstream toolkits expose a single integer, n_top_genes, that is usually left at a value near 2,000^4^. Two distinct decisions are conflated within that integer, and neither is well specified. The first is list length: how many genes are necessary to resolve the true structure, recognizing that too short a list erases biological signal, while too long a list adds noise and computational cost. The second is cutoff fairness: because global ranking measures variability across all cells, markers for rare cell types often lose the “variance vote count” to bulk tissue variation. Consequently, informative rare markers frequently land just below a hard top-k cutoff, leading to an unfair allocation of features when abundant populations dominate the matrix.

Large-scale benchmarks show that no single HVG formula wins everywhere and that mixtures of diverse scores can be more robust than any one of them ^5–7^. Difficulty-aware evaluations further show that easy cell-type tasks tolerate weak gene sets, whereas subtle structure is sensitive to both list length and selection strategy ^8,9^. Changing the ranking formula and changing the list length are therefore distinct levers, and routine computational pipelines still need a defensible default for cardinality on top of a familiar ranking—in practice, seurat_v3 via Scanpy ^3,4,10^.

Automatic list length is not itself novel. Threshold-based selectors already exist: getTopHVGs in scran can return genes by an FDR or residual-variance threshold rather than a fixed integer ^11^; M3Drop applies a dropout-model FDR ^12^; deviance-based selection replaces the variance statistic entirely ^13^; and neighborhood-based scores such as triku avoid a single global *n* ^14^. Elbow heuristics, library-size rules, and tissue lookups can also adjust *n*. However, all of these either change the selection criterion (e.g., applying a statistical threshold on a per-gene score) or use metadata not derived from the embedding geometry of the matrix at hand. What remains missing is a ranking-agnostic, Scanpy-native default—one that holds a user-chosen global ranking fixed while dynamically setting *k* based on the multi-seed density and stability features of that specific matrix, incorporating conservative floors back to 2,000 when geometric evidence is weak (Supplementary Table S2). Our primary claim is therefore focused on this design axis—cardinality under a frozen ranking—rather than universal clustering gains.

Here we describe scFair, an HVG layer built on a single premise: the ranking is the user’s choice and should be left alone, while the truncation point is a modeling decision best derived from the data (Figure 1). Geometry-aware auto_n reads multi-seed density and stability structure from an intermediate embedding and returns a base size k, operating on the principle that the extra compute is a worthwhile trade-off for a safer, data-driven default. A same-rank append step then extends the chosen list by a short near-miss tail. Rather than introducing aggressive cluster-conditional reallocation, this append step serves as a conservative safeguard against cutoff unfairness, ensuring that informative genes residing just below the hard threshold are not arbitrarily discarded.

**Figure 1.**
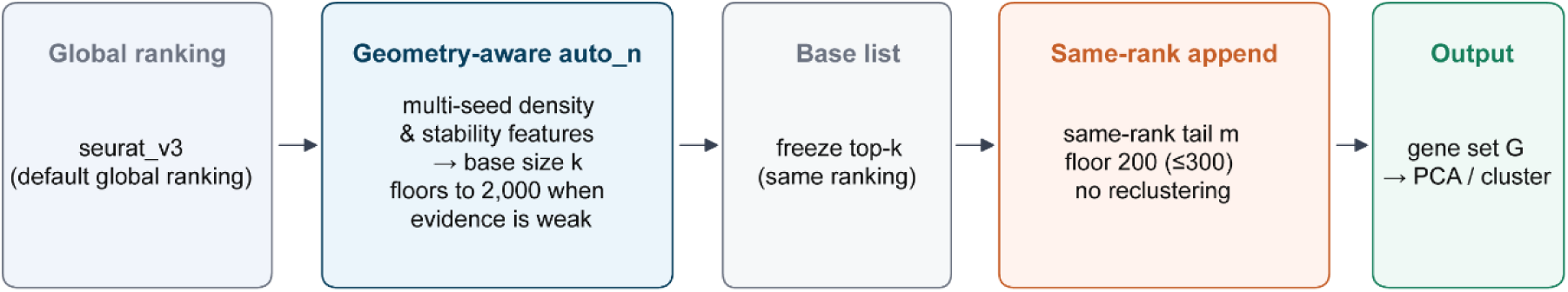
The scFair selection path. A standard global ranking is computed first and is never modified. Geometry-aware auto_n sets the base list size *k* from multi-seed density and stability features (near 2,000 by default; shorter or longer only with geometric support). Same-rank append adds a near-miss tail *m*. Both components ship together.

We evaluate end-to-end Leiden clustering agreement (ARI and NMI ^15^) against curated cell_type labels on a unified panel of 18 datasets. Oracle *k*-sweeps first establish the available optimization margin within the cardinality axis under a fixed ranking. The proposed pipeline is then measured against the classical default and a longer fixed default, decomposed into its two arms, checked on a Tabula Sapiens organ holdout ^16^, and compared with three classes of alternatives: fixed-*k* rank fusion (which varies content instead of length); cell-number-only and FDR-threshold length rules (which vary length by other means); and the neighborhood-based selector triku (which varies both; Section 2.8). A leave-one-dataset-out re-selection of the short-list thresholds and a multi-metric secondary suite bound how much of the result could be attributed to calibration artifacts.

## 2. Results

### 2.1 Evaluation design

Primary claims use a unified panel of 18 labeled datasets assembled from public benchmarks and atlases (sources in Section 5; key references ^7,16–26^), stratified by label provenance into GOLD-13 (orthogonal or high-quality labels) and MIXED-5 (author or multi-protocol labels) and reported as such (Table 1; Figure 5A). Three layers of evidence are kept distinct throughout. Oracle *k*-sweeps under seurat_v3 measure the size of the cardinality axis: best_n = 2,000 on only 1/18 datasets, with mean ARI(best_n) − ARI(2000) = +0.033 (median +0.026). This is an upper bound obtained by selecting *k* on the evaluation metric, so it bounds what any length rule could achieve rather than what one should expect. Panel-18 is the primary benchmark for the shipped path against HVG@2000; because rule thresholds were iterated with partial overlap to this panel (Methods 3.3), it is a calibrated benchmark (Section 2.11 quantifies the exposure). Test-5, an independent organ set assembled after the rule was frozen (Section 2.9), provides a directional external check.

**Table 1.** Evaluation panels.

| Panel | *n* | Role | ΔARI vs HVG@2000 |
| --- | --- | --- | --- |
| Panel-18 (unified) | 18 | Primary calibrated benchmark | median +0.016; 13/5; $P = 0.0077$ |
| GOLD-13 | 13 | Orthogonal / high-quality labels | mean +0.0118; 9/4 |
| MIXED-5 | 5 | Author or multi-protocol labels | mean +0.0247; 4/1 |
| Test-5 (Tabula Sapiens organs) | 5 | Independent holdout; not used for tuning | median +0.0131 (IQR $-0.0003$ to $+0.0154$); 3/2; $P = 0.31$ |

Unless stated otherwise, ranking is seurat_v3 on raw counts; the shipped path is n_top_genes=“auto”, balance_method=“append”, mitochondrial and ribosomal filters off, and automatic Leiden resolution drawn from {0.8, 1.5}. Baselines are scanpy HVG@2000 and HVG@2200. The primary metric is mean ARI over two Leiden seeds. Nonparametric tests are two-sided Wilcoxon signed-rank tests on paired per-dataset means; bootstrap percentile intervals for the median use 10,000 dataset resamples. Throughout, paired sign counts are written positive/negative, or positive/negative/exactly zero where ties occur; counts written as “x/18 datasets” are counts out of the full panel. The pre-specified primary contrast is the shipped path versus HVG@2000 on panel-18; all other contrasts, including the strata of Section 2.4, are secondary or post hoc and are interpreted without family-wise error control.

### 2.2 Design of the selection path

The shipped path is summarized in Figure 1. scFair computes a standard global ranking first and never modifies it: no new variance model is introduced, no gene is reweighted by cell_type, and no intermediate clustering feeds back into the score. Everything downstream operates on the rank order as given, which is what makes cardinality separable from ranking content and lets the same layer sit on top of any flavor the user prefers.

The quantity auto_n must estimate is how many genes are needed to resolve the structure that is present, and the informative signal for that is not cell count but how the cells are arranged. We therefore extract features from the embedding itself rather than from metadata. For each of three seeds, an intermediate 2,000-gene seurat_v3 pipeline is run to a neighbor graph and a Leiden partition, and two families of statistics are read off: density geometry in a low-dimensional embedding — the depth *v* of the valleys separating density modes, the fraction *f* of valleys that are shallow, and the number n_d of density cores — and partition stability across bootstraps. The rationale is direct. Deep valleys around many cores mean the populations are already separated, so a short list suffices and a long one mainly adds variance-dominant genes from abundant populations. Many shallow valleys around few cores mean structure is present but unresolved, which is the regime where additional genes buy resolution. Low stability across seeds means the geometry itself is not trustworthy, which must not be read as evidence for either extreme. Because a single seed can be misleading, features are aggregated by their median across seeds and the confidence flag is taken from the worst seed.

These statistics enter an ordered, first-match-wins decision list of nine conditions in falling-rule-list form (Methods 3.3; Supplementary Table S1a). Only the long regime is continuous, scaling *k* with stability between 2,500 and k_max; the remaining rows assign one of four fixed sizes. The decision list is then filtered by six confidence guards applied in fixed order (Supplementary Table S1b), and the guards are where the design philosophy lives: a default ladder buffer lifts short and mid raw sizes one rung toward 2,000 unless the geometry indicates a well-resolved short list, any budget obtained under low structural confidence is floored to 2,000, and a residual guard catches short lists resting on untrusted density estimates. The intended operating point is a high-specificity, low-sensitivity deviation detector: on most matrices auto_n returns, or is floored to, approximately 2,000, and it departs from that value only when multi-seed geometry supports the move. The cost of this choice is deliberate and visible in the results below — the rule declines many opportunities the oracle sweep identifies — and the benefit is that a wrong short list, which is the failure mode that damages a downstream analysis most, is rare.

The second component, append, freezes the top-*k* list and adds a short same-rank tail *m*. It never removes a gene from the base list, never reclusters, and never consults the geometry for gene identity, only for the length of the tail. It exists because the cutoff rank is not a meaningful boundary in a continuous score: genes immediately below it are near-indistinguishable from those immediately above, and a hard truncation discards them for no statistical reason. We evaluate auto_n and append jointly as the shipped path. The two arms act on disjoint sets of datasets and are not presented as equally well-supported mechanisms (Section 2.4).

### 2.3 End-to-end gains on panel-18

Selected list lengths *n* = *k* + *m* span short regimes (*k* ≈ 500), one moderate contraction (*k* ≈ 1,750), and long mixtures (*k* ≈ 3,000–4,400), with many datasets remaining near 2,000 (Figure 2A; full budgets in Supplementary Figure S1). Against HVG@2000, the shipped path yields median ΔARI = +0.0160 (mean +0.0154; SD 0.022; IQR [−0.0015, +0.0228]; 13/5; *P* = 0.0077; bootstrap 95% CI for the median [+0.0009, +0.0226]; Figure 2B), or roughly half of the mean oracle headroom available under the same ranking. The gain is on partition agreement rather than embedding geometry: mean ΔNMI = +0.0065 (15/3; *P* = 0.0066), while silhouette width, *k*-NN accuracy, and LISI are flat (Supplementary Note S1, Supplementary Figure S6). This matters for interpretation, because a gene set can tighten an embedding without changing the partition recovered from it; here the improvement is on the partition itself. Arm decomposition follows in Section 2.4; the independent holdout is reported in Section 2.9.

**Figure 2.**
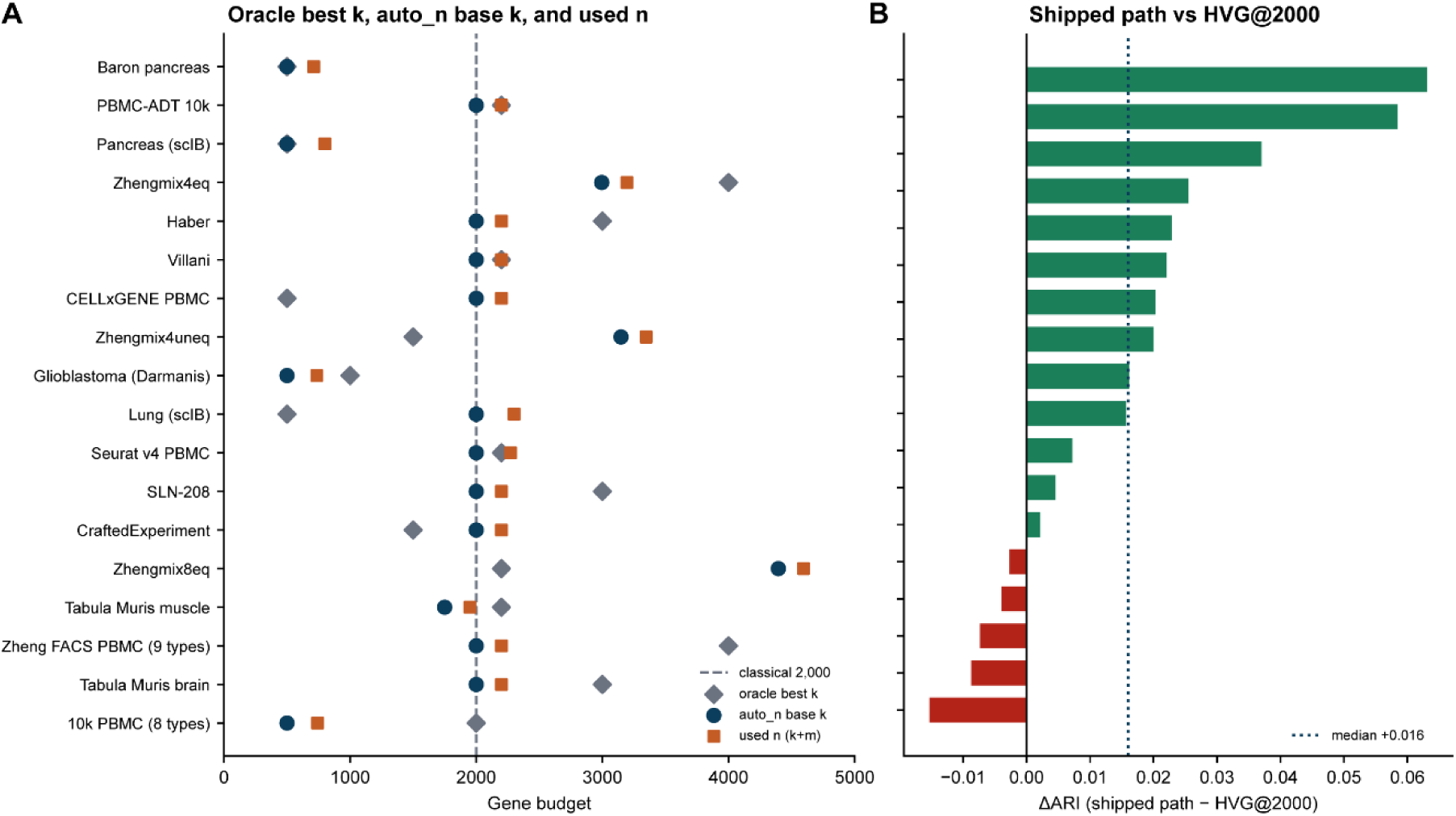
Gene budgets and end-to-end effect on panel-18. (**A**) Oracle best *k*, auto_n base *k*, and used *n* after append, per dataset. (**B**) Paired ΔARI for the shipped path versus scanpy HVG@2000 (dotted line, median).

### 2.4 Arm decomposition and post hoc *k*-move strata

To separate the two components we compared HVG@2000, sizing-only (automatic *k* without append), and the shipped path on panel-18 (Figure 3). Marginally, sizing-only has median ΔARI ≈ 0, with exact zeros on the 10 datasets where *k* = 2,000 and 7/1 among the 8 datasets where *k* moves (full-panel mean +0.008; *P* = 0.11). Append versus sizing-only has median +0.0068 (12/6; *P* = 0.18). Only the combined path is significant (median +0.016; 13/5; *P* = 0.0077). Neither component reaches panel-wide significance alone, which is expected when each acts on a subset of the panel and is zero elsewhere.

**Figure 3.**
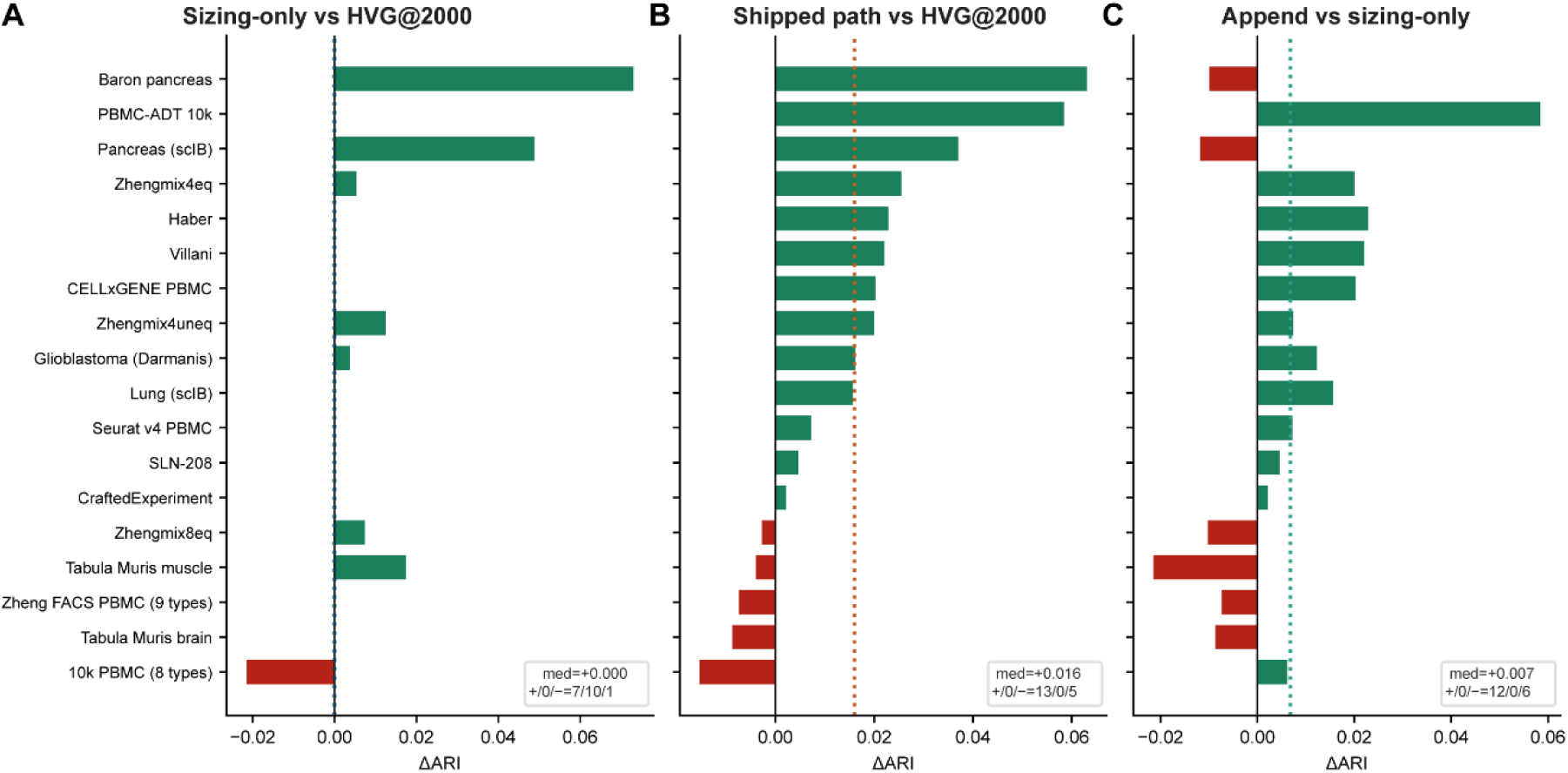
Arm decomposition on panel-18. (**A**) sizing-only versus HVG@2000. (**B**) Shipped path versus HVG@2000. (**C**) Append versus sizing-only. Positive deltas in green, negative in red; dotted line, median.

Stratifying on whether auto_n moves *k* (|*k* − 2,000| > 50) resolves the pattern (Figure 4; per-dataset sizing versus append in Supplementary Figure S2). This split was defined after inspecting the results and the strata are small (8 movers, 10 stayers), so we report stratum medians and sign counts without confirmatory subgroup *P* values.

**Figure 4.**
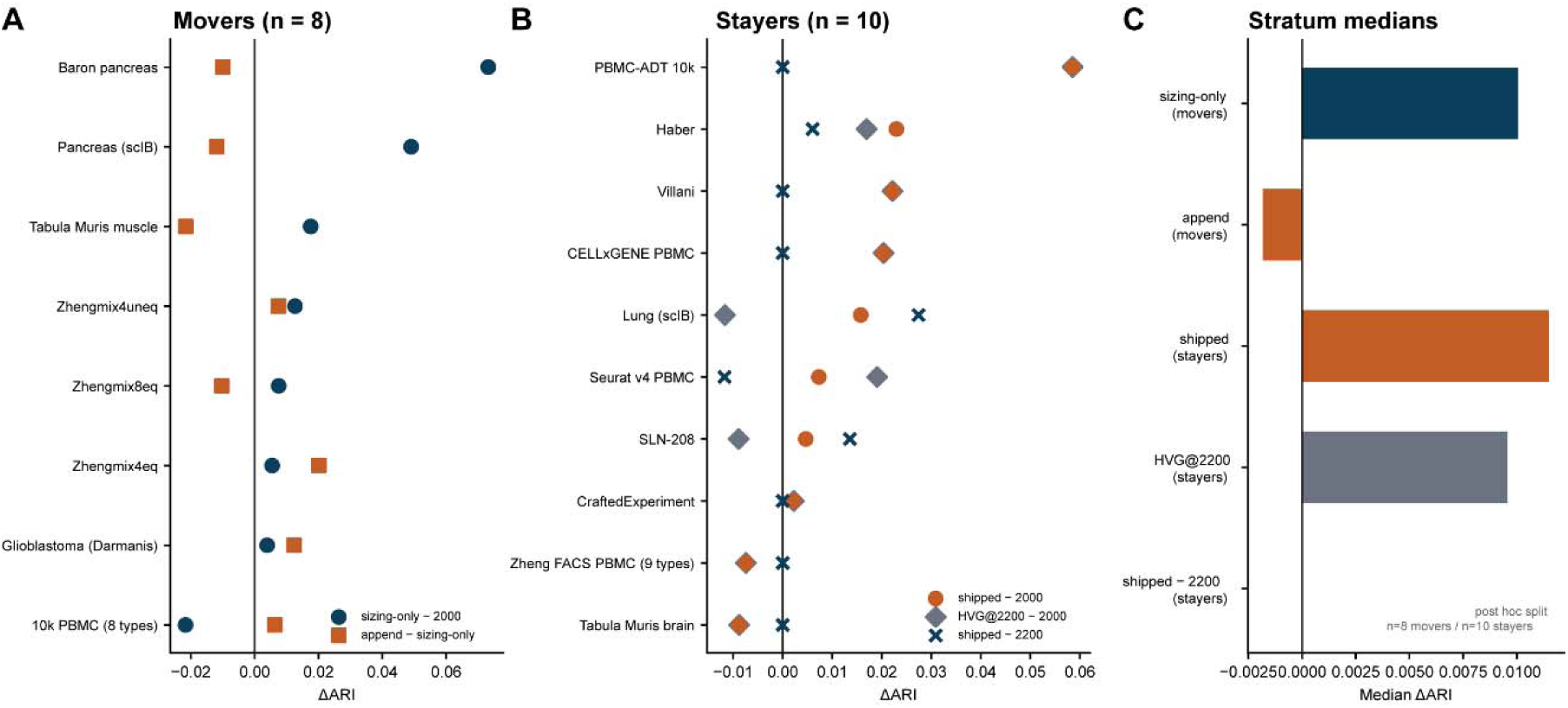
*k*-move strata on panel-18 (post hoc split; Section 2.4). (**A**) Movers (*n* = 8): sizing-only − HVG@2000 alongside append − sizing-only. (**B**) Stayers (*n* = 10): shipped − HVG@2000, HVG@2200 − HVG@2000, and shipped − HVG@2200. (**C**) Stratum median ΔARI. On *m* = 200 stayers the shipped path coincides with HVG@2200 except where a score tie at the cutoff rank is broken differently (Methods 3.4).

*Movers (n = 8).* Sizing carries the effect: sizing-only median ΔARI = +0.010 (mean +0.018; 7/1), with large gains on compact pancreas matrices (Baron pancreas +0.073; multi-protocol human pancreas (scIB) +0.049) and one spurious short-list loss (10k PBMC (8 author types), −0.022). Append is near neutral on this stratum (median append − sizing-only = −0.002; 4/4).

*Stayers (n = 10).* sizing-only is identically zero by construction, so the shipped path reduces to append. On 8/10 of these datasets *m* = 200, and the remaining two receive *m* = 272 and *m* = 300 after a density-driven increase, so in most cases the selected set is the seurat_v3 top-2200 list. Empirically, median (shipped path − HVG@2200) = 0, with exact zeros on 6/10; the other two *m* = 200 stayers (Haber, SLN-208) diverge from a literal top-2200 list by a handful of genes because the seurat_v3 score has an exact tie at the rank-2200 cutoff and scFair’s rank construction is not coordinated with scanpy’s native tie-break (Methods 3.4). Median (HVG@2200 − HVG@2000) = +0.010. The stayer “append effect” is therefore automatic adoption of a fixed 2,200-gene list, not a second geometric mechanism.

Two-sided Mann–Whitney and permutation tests that append helps more on stayers than movers are borderline (*P* = 0.083 and 0.088), but the direction is instructive: append is a proportionally larger perturbation on movers (+200 genes on a base of 500, +40%) than on stayers (+10% on 2,000), yet its apparent benefit concentrates on stayers. That is what one expects if the stayer effect is the 2,000 → 2,200 step itself while movers have already realized their gain through *k*. The panel-wide result is therefore a mixture of two arms with unequal evidential support: automatic list lengthening acting on the majority, and geometric resizing acting on a minority that is smaller, post hoc stratified, and examined further in Section 2.5.

### 2.5 Is a fixed HVG@2200 default a sufficient substitute?

Because the shipped path equals seurat_v3 top-2200 on most stayers, the natural challenge is whether users should simply raise the global default to 2,200. Across the full panel, HVG@2200 versus HVG@2000 is not significant (median ΔARI = +0.0045; 11/7; *P* = 0.17), but that contrast averages over strata that behave differently. On stayers (*n* = 10), median (HVG@2200 − HVG@2000) = +0.010 and median (shipped − HVG@2200) = 0, as set identity requires. On movers (*n* = 8), median (HVG@2200 − HVG@2000) = +0.003 (5/3), so there is no free lift from lengthening, while median (shipped − HVG@2200) = +0.007 (5/3) (Figure 5B; Supplementary Figure S5). Lengthening and resizing are therefore not interchangeable: the mover stratum contains both lengthened and shortened lists, and one fixed value cannot approximate both.

**Figure 5.**
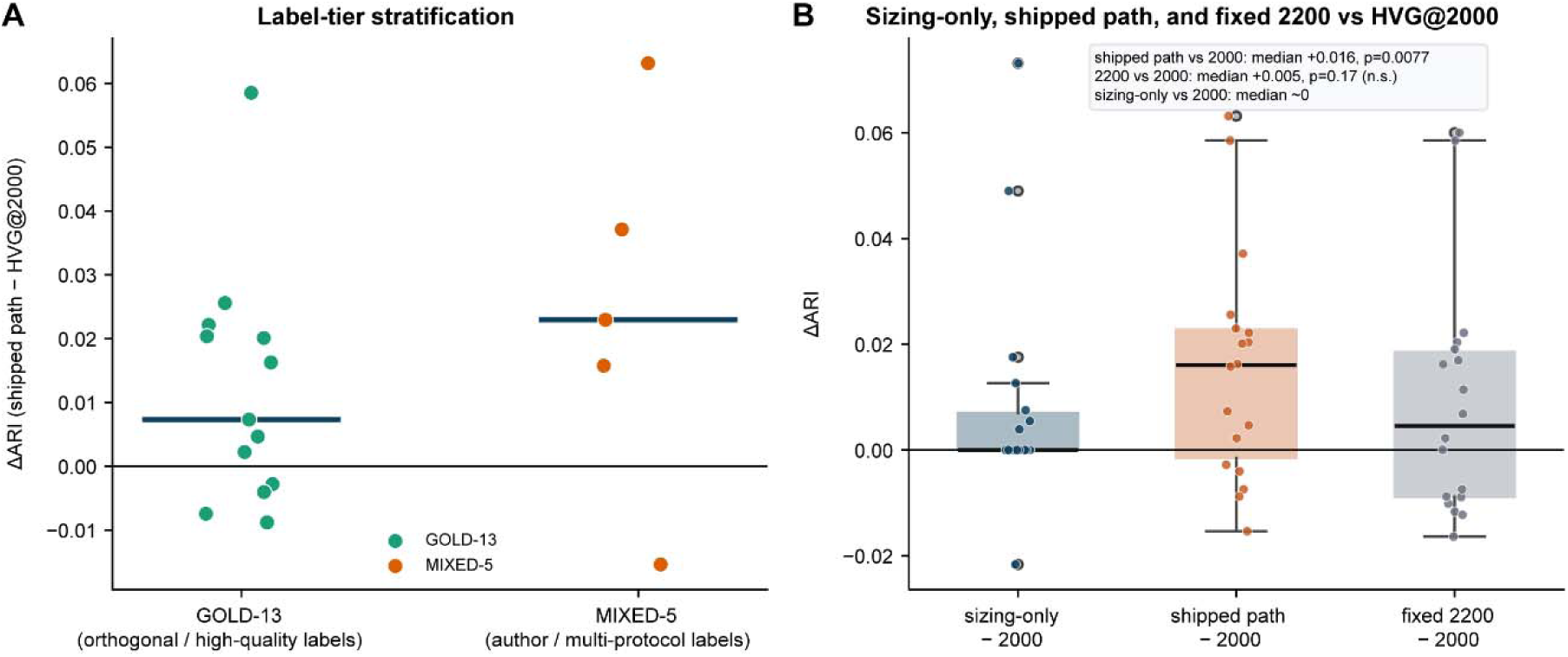
Label-tier and HVG@2200 controls on panel-18. (**A**) Shipped-path ΔARI stratified by GOLD-13 versus MIXED-5 label tier (Section 2.1); horizontal bars, stratum medians (Table 1 reports stratum means). (**B**) Distribution of ΔARI for sizing-only, the shipped path, and fixed HVG@2200, each relative to HVG@2000 (Section 2.5).

The second case is the sharper test. Restricting to short-*k* movers (sizing-only *k* = 500; *n* = 4, post hoc) gives median (shipped − HVG@2200) = +0.029; a default of 2,200 cannot produce a 500-gene list, and no single fixed value can serve both these datasets and the rest of the panel. These four datasets are not exchangeable, however: Baron pancreas (+0.063) and multi-protocol human pancreas (scIB) (+0.054) dominate, glioblastoma (Darmanis) is weakly positive (+0.005), and 10k PBMC (8 author types) is negative (−0.003 against HVG@2200; −0.015 against HVG@2000), a spurious short list. We regard the short-list branch as the mechanistic core of the method and, on the present evidence, as its narrowest claim: it is carried by two pancreas objects plus one weak non-pancreas case, and it is not confirmed on the holdout (Section 2.9). Baron pancreas warrants caution for a second reason — it is one of only three datasets on which an external content-based selector outperforms the shipped path (Section 2.8).

### 2.6 Same-rank append

Global feature ranking measures variance across an entire dataset, inherently skewing toward genes highly expressed in abundant populations. As a result, critical markers for rare cell types often accumulate just below a hard top-k threshold, losing the variance vote count to bulk tissue variation. This creates an unfair allocation of features in standard fixed-length lists. Rather than introducing cluster-conditional reweighting or per-cluster quotas, the append mechanism (balance_method=“append”) provides a conservative safeguard against this cutoff unfairness. It freezes the primary top-k list B and adds a short same-rank tail of near-miss genes, returning G = B ⍰ {next m genes in the same ranking}. The tail length m is floored at 200, with an optional density-driven extension to 300. The procedure never drops genes from B and requires no intermediate reclustering. When k = 2,000 and m = 200, G reproduces the seurat_v3 top-2200 list, subject only to minor tie-break discrepancies at the cutoff (Methods 3.4). This condition applies to 8 of 10 panel-18 stayers. Append operates as a residual mechanism—securing a slightly longer fixed list when geometry leaves k at 2,000, and serving as a small additive tail otherwise. As a marginal full-panel contrast, its independent contribution is not statistically significant (median +0.0068; P = 0.18). Users requiring a strict gene budget should set balance_method=“none”.

### 2.7 Ranking content at fixed *k*

Holding *k* at the auto_n value isolates the other lever. Mean-rank fusion of seurat_v3, seurat, cell_ranger, and pearson_residuals substantially changes list membership (Jaccard commonly 0.3–0.7; Supplementary Figure S4) but does not improve clustering robustly: panel-18 median ΔARI = +0.015 (mean +0.0035; SD 0.029; IQR [−0.006, +0.020]; 13/5; minimum −0.086; 10th percentile −0.04; *P* = 0.32); Test-5 median = −0.009 (2/3; *P* = 0.44; Figure 6). The fusion median is close to the shipped path’s +0.016, so the difference is dispersion and left-tail risk rather than central tendency (shipped path SD 0.022, minimum −0.015); these are descriptive tail summaries, not a formal test. We do not adopt ensemble ranking as a default: a rule whose median matches the shipped path but whose mean is four-fold lower, and which occasionally loses 0.08 ARI, is a poor default. Ranking content and list cardinality remain separable levers, and score-oriented methods stay relevant when the ranking, not *k*, is the failure mode.

**Figure 6.**
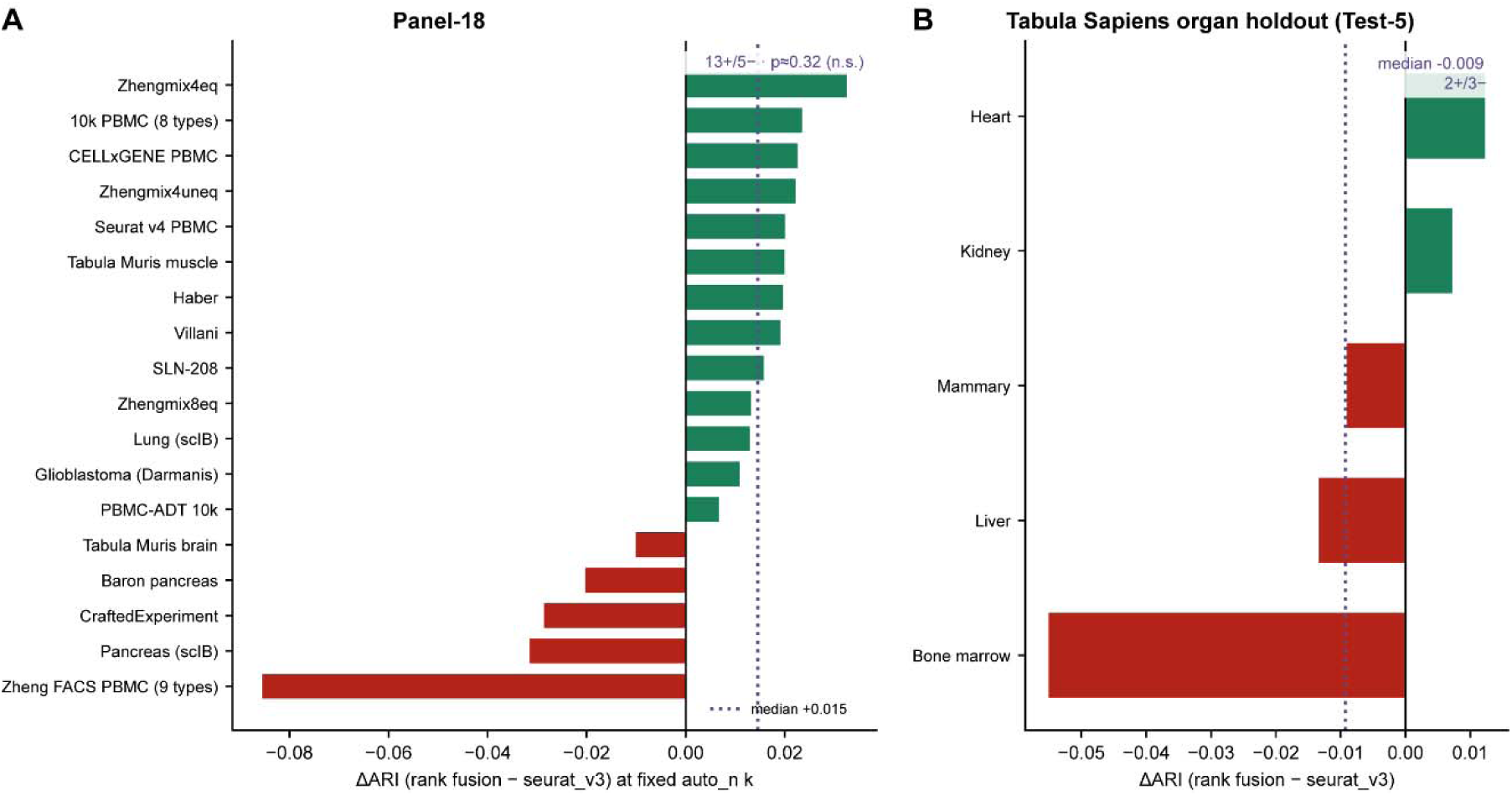
Fixed-*k* multi-method rank fusion versus seurat_v3. (**A**) Panel-18. (**B**) Tabula Sapiens organ holdout (Test-5).

### 2.8 End-to-end comparison with an external selector (triku)

The preceding ablation varies ranking content within our own framework. We also compared against an external selector that acts on content directly. triku ranks genes by localized *k*-nearest-neighbor expression — the Wasserstein distance between a gene’s observed *k*-NN expression distribution and a random-neighbor null — and can choose list length automatically ^14^. It therefore operates primarily on gene content and only secondarily on *n*, whereas scFair freezes the seurat_v3 rank order and automates cardinality alone. The comparison is consequently end-to-end rather than a single-lever ablation, but it answers the practical question of which default selection path better supports Leiden clustering under a shared protocol.

#### Protocol

We used triku 2.2.0 from PyPI with author defaults for all scoring hyperparameters (n_features=None for automatic length, s = −0.01, n_windows=75, min_knn=6, dist_conn=“dist”, distance_correction=“median”); no grid search or dataset-specific tuning was performed. The package requires a neighbor graph, so following the documented scanpy workflow we applied filter_genes(min_cells=5), normalize_total(10□), log1p, PCA (≤ 50 components), and neighbors (15 neighbors, ≤ 30 PCs) before calling tk.tl.triku(…, use_raw=False). Setting use_raw=False was required because log-normalized expression was placed in .X, whereas the package default expects counts in .raw. Downstream evaluation was identical across all arms (Section 3.5), with two Leiden seeds.

Against the classical default, untuned triku underperformed HVG@2000 on 13/18 datasets (median ΔARI = −0.018; mean −0.025; IQR [−0.033, +0.007]; *P* = 0.054), so the deficit is not an artifact of comparing only against scFair. Relative to the shipped path, triku was worse on 15/18 datasets (median ΔARI for shipped − triku = +0.024; mean +0.040; IQR [+0.008, +0.067]; *P* = 0.0040; Figure 7A). triku returned shorter lists overall (median *n* ≈ 1,522; range 644–2,964) than the shipped path (median 2,200; range 712– 4,595; Figure 7B), and won on three datasets: Baron pancreas, Zhengmix8eq, and Tabula Muris brain.

**Figure 7.**
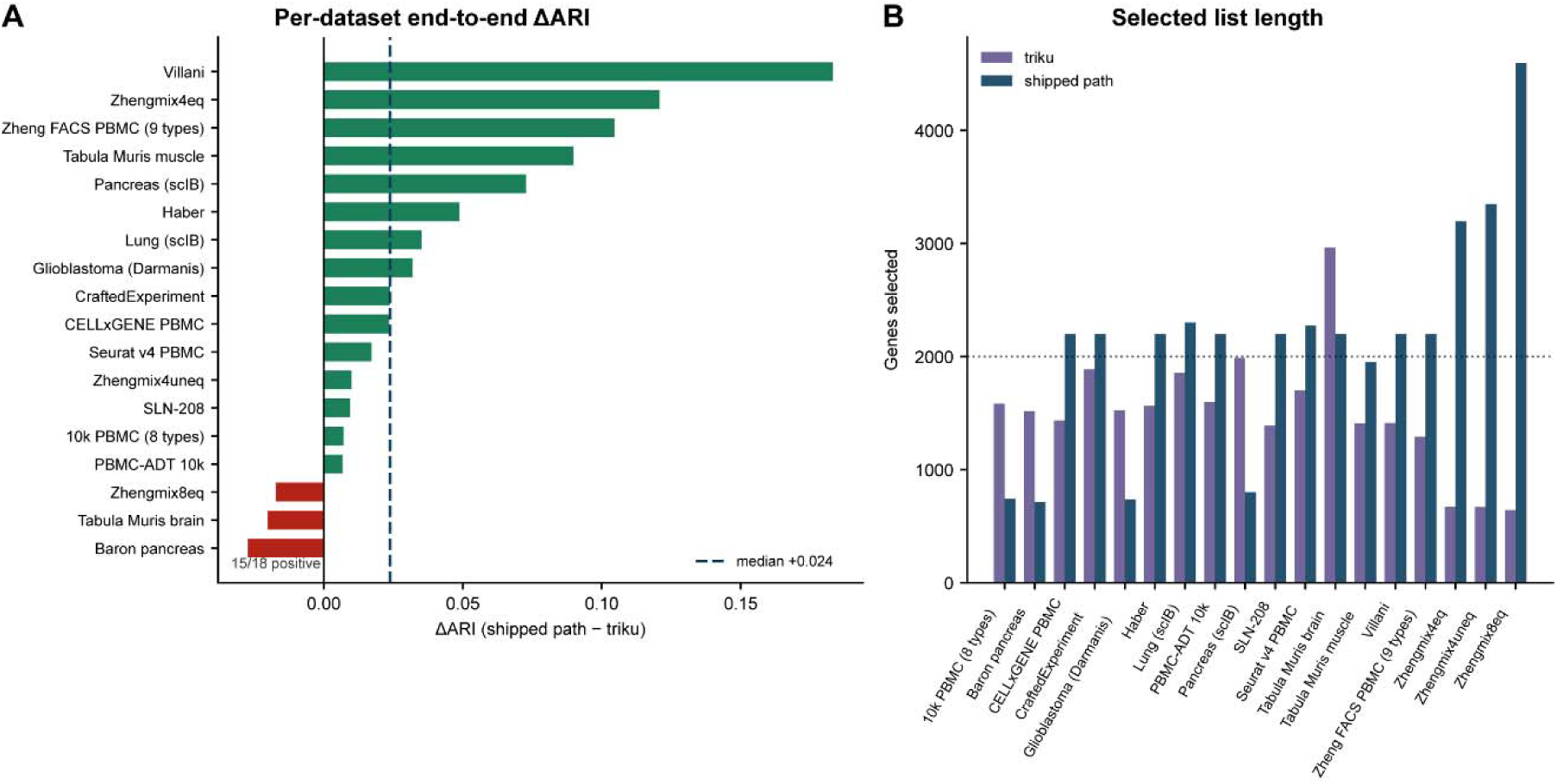
Shipped path versus triku on panel-18 (two seeds; triku 2.2.0, author defaults, no tuning). (**A**) Per-dataset ΔARI (shipped − triku); dashed line, median (+0.024; 15/18 datasets). (**B**) Selected gene counts per method (dotted line, 2,000). For reference, triku − HVG@2000 has median ΔARI = −0.018 (13/18 datasets).

Two caveats bound the interpretation. Because triku changes both which genes enter the list and how many, this is an end-to-end comparison of default paths rather than a length-only control; unlike the FDR baseline of Section 2.11, we do not freeze seurat_v3 at triku’s chosen *n*, so content and cardinality remain confounded by design. And one of triku’s three wins is Baron pancreas, which is also one of the two datasets driving the short-*k* advantage of Section 2.5, so a content-based selector outperforms geometric resizing on the very matrix where resizing looks strongest. Within these limits, clustering agreement on panel-18 favors the shipped path, and default triku does not improve on classical HVG@2000.

### 2.9 Independent holdout: Tabula Sapiens organs

Test-5 comprises five organs (liver, heart, bone marrow, kidney, mammary) from Tabula Sapiens 10x 3′ v3 with manually curated Cell Ontology labels ^16^. These objects were assembled after the structure rule was frozen and were not used to adjust thresholds, branch order, or floors. The protocol matches panel-18.

The shipped path versus HVG@2000 gives median ΔARI = +0.0131 (mean +0.0114; IQR [−0.0003, +0.0154]; 3/2; *P* = 0.31). sizing-only versus HVG@2000 gives median 0: four of five organs receive *k* = 2,000, by default or by a fine-mode floor, and heart alone retains a short base (*k* = 500, ΔARI = +0.0027). Append versus sizing-only gives median +0.0128 (3/2; *P* = 0.31) (Supplementary Figure S3).

With five organs the holdout cannot support panel-level significance, but it is informative about which branch transfers. On the four *k* = 2,000 organs the shipped path tracks append — the automatic 2,000 → 2,200 extension — exactly as the stayer analysis predicts. The mover branch, which is where the path differentiates itself from a fixed 2,200 default, is not confirmed: the single mover shows a marginal sizing benefit rather than a pancreas-scale short-*k* gain. The holdout thus reproduces the arm that a simpler rule could also deliver and leaves the distinctive arm untested.

### 2.10 Cell-number-only cardinality baselines

A cheaper alternative to density-based sizing is a rule that uses only cell count. Under the same ranking, seeds, and clustering protocol we compared HVG@2000, sizing-only, and two a priori *n*_obs schedules that were not fit to panel-18 ARI: (i) a step table (*n* < 2,000 → 1,000; < 5,000 → 1,500; < 10,000 → 2,000; < 20,000 → 2,500; otherwise 3,000), and (ii) *k* = clip(round(20√*n*), 500, 4,000).

Neither rule beat HVG@2000 (step: median ΔARI = 0, mean −0.007, 7 positive / 8 negative / 3 zero, *P* = 0.30; √*n*: median ≈ 0, mean −0.009, 9/9/0, *P* = 0.47). sizing-only versus the step table was directionally positive (median +0.006, mean +0.016, 9/8/1, *P* = 0.13) as was sizing-only versus √*n* (median +0.007, mean +0.017, 11/7/0, *P* = 0.18). The mechanism behind the largest differences is explicit: on large but compact pancreas matrices auto_n selects short lists (*k* = 500) whereas cell-count tables lengthen the list by construction (multi-protocol human pancreas (scIB), +0.101 versus the step table; Baron pancreas, +0.073). Where auto_n leaves *k* at 2,000, cell-count rules occasionally edge ahead simply by landing on a different fixed length. Cell-number tables therefore neither beat the classical default nor recover the short-*k* compact regimes that anti-correlate with cell count.

### 2.11 FDR-threshold length baseline and leave-one-out re-calibration

A score threshold is the closest classical route to automatic length. As a length-rule baseline in that class — not a package-faithful Bioconductor re-run of scran getTopHVGs — we used a Python residual-variance selector with Benjamini–Hochberg control at q = 0.05, which yields median *n* = 2,282 (range 1,479–3,361; the [200, 5,000] clip is never reached). Using that rule’s own gene content gives median ΔARI = −0.018 versus HVG@2000 (4/14; *P* = 0.13), worse than the fixed default despite a comparable length.

Because residual ranking differs from seurat_v3, that contrast confounds content with length. The clean cardinality control freezes seurat_v3 and takes top-*n* at the FDR-chosen length: median ΔARI versus HVG@2000 = +0.0005 (10/8; *P* = 0.64), essentially flat, and a median +0.025 above the residual gene content at the same *n*. Two conclusions follow, and they are the reason we treat how *k* is chosen as the substantive claim rather than the fact that it is chosen automatically: automatic length under a fixed ranking does not beat 2,000 by itself, and the harm from the FDR selector was almost entirely ranking content, consistent with Section 2.7. sizing-only versus FDR-length-with-seurat_v3 is likewise flat across the full panel (median ≈ 0; *P* = 0.64), as expected when most datasets stay at *k* = 2,000; the short-*k* movers of Section 2.5 remain the only locus where geometric sizing separates from length rules.

To test local over-fitting of the short-list thresholds, we re-selected the two short-hard cutoffs by leave-one-dataset-out over an 81-point vm/nd grid, using oracle seurat_v3 *k*-grid ARI on the 17 training datasets and evaluating the held-out one. No fold (0/18) selected constants different from the shipped defaults, so held-out ΔARI equaled the default throughout. Two readings are compatible and we state both: the constants are stable under single-dataset leave-out within this grid, and 0/18 is also what one expects if those constants were already near ARI-optimal on this panel, so the analysis cannot refute calibration overlap. The test is also partial, since only 9/18 held-out datasets enter a short-hard branch in the default trace (4 remain at *k* = 500; 5 are buffered or floored to 2,000) and the confidence guards of Supplementary Table S1b, which determine most classical-2,000 outcomes, were not re-tuned. We therefore claim stability only within the short-hard threshold neighborhood, not full-rule out-of-sample validity.

## 3. Methods

### 3.1 Overview

scFair is a preprocessing layer for AnnData objects. Its interface is a global HVG ranking, an automatic or fixed base *k*, and an optional same-rank extension. The design principles are: keep the ranking standard and frozen; make cardinality the only automated default; keep secondary mechanisms minimal; and avoid cluster-dependent re-ranking in the default path.

### 3.2 Global ranking

The default flavor is seurat_v3 on raw counts ^3,4,10^. Optional arguments include batch_key, mitochondrial and ribosomal filters (off by default), and marker_genes with marker_mode=“force” (default “none”) for user-directed inclusion of named genes; the marker path is a manual rescue and is not part of the evaluated default (Limitations).

### 3.3 auto_n

With n_top_genes=“auto”, the base size *k* is chosen by the structure-rule implementation (shipped version 7). The procedure is deterministic given random_state. Neither the ranking nor the feature extraction reweights genes by cell_type. Optional type labels, when provided, enter exactly one post-rule guard: the true short-list exception, which retains *k* = 500 when short-hard geometry co-occurs with n_d ≥ 10 and n_types ≥ 5. Without labels that guard cannot fire and the soft buffer toward 2,000 applies instead. On panel-18 runs with type labels supplied, this guard appeared in 6/18 rule traces, of which 4 retained *k* = 500 and 2 were subsequently floored to 2,000 in fine mode.

#### Feature extraction

For each seed *s* = 0, 1, 2 (random_state + *s*), an intermediate seurat_v3 list of 2,000 genes on counts is followed by normalize_total(10□) → log1p → scaling → PCA → neighbor graph → Leiden at resolution 0.5 ^15^. This pipeline yields bootstrap pair-stability over 5 bootstraps (mean stability μ_s and worst-seed stability μ_s,min) and 3D density-valley statistics on the embedding: valley depth *v* ∈ [0,1], the fraction of shallow valleys *f*, the number of density cores *n*_d, and a density-confidence flag κ ∈ {high, medium, low}. Per-seed features are aggregated by their median, with confidence taken from the worst seed.

#### Base cardinality

The aggregated statistics are mapped to a raw base size *k*□ through an ordered, first-match-wins decision list R = [(c□,*k*□), …, (c□,*k*□)] in the falling-rule-list form of Wang and Rudin ^27^. The ordering encodes the geometric argument of Section 2.2: sharply separated geometry with many resolved cores (high *v*, high n_d) is assigned a short list; diffuse geometry with many shallow valleys and few cores (high *f*, low n_d) is assigned a long list scaled continuously with stability; intermediate valley-depth and core-count combinations map to fixed mid-range sizes; and low-signal or ambiguous cases fall through to the classical default. Only the long-regime assignment is continuous:

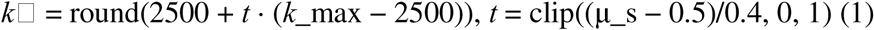

The remaining eight conditions and their exact thresholds are given in Supplementary Table S1a. Confidence guards are then applied in the order listed in Supplementary Table S1b. A default one-rung buffer lifts short and mid raw sizes toward 2,000 unless the geometry indicates a well-resolved short list, and any base size obtained under low structural confidence is floored to 2,000. Five of the six guards move *k* toward the classical default; the sixth, the true short-list exception, only withholds that correction when labeled types corroborate short-hard geometry. No guard moves *k* further from 2,000, which is the formal statement of the high-specificity operating point described in Section 2.2. Diagnostics — regime, guard trace, and per-seed *k* — are recorded in the AnnData object. Seeds are combined as follows: if all three per-seed values of *k* agree, that value is used; otherwise *k* is recomputed from the aggregated features, and for n_obs ≥ 10,000 a short-regime vote (*k* ≤ 500) from a single seed cannot override an aggregated mid- or long-regime result (*k* ≥ 1,500). The source of truth for all constants is the structure-rule implementation (shipped version 7).

#### Calibration

Thresholds and branch order were iterated during method development on datasets that substantially overlap panel-18, with feature diagnostics and, in later rounds, ARI feedback. Leave-one-dataset-out re-selection over the short-hard vm/nd constants (Section 2.11) never selects non-default values (0/18 folds), which is compatible both with local stability and with constants already fit to this panel; only 9/18 datasets exercise a short-hard branch, and the Supplementary Table S1b guards were not re-tuned. We do not claim a prospectively registered rule.

### 3.4 Append

With balance_method=“append”, the returned set is *G* = top-*k* ∪ {next *m* genes in the same ranking}, with *m* ≥ 200 and an optional density-driven increase to at most 300. When the global score has an exact tie at the cutoff rank, the specific genes admitted can differ by a few from a freshly called scanpy.pp.highly_variable_genes at the same list length, because scFair’s rank construction (pandas rank, average method) and scanpy’s internal tie-break (double argsort) are not coordinated; this affected 4/18 panel-18 datasets — Haber and SLN-208 at the rank-2200 cutoff (stayers), and Zhengmix8eq and Tabula Muris muscle at their own selected-length cutoffs (movers) — by 2–100 genes each (Sections 2.4, 2.6). balance_method=“none” returns exactly the top-*k* list.

### 3.5 Downstream evaluation

Each selected gene set was processed identically: subsetting → normalize_total(10□) → log1p → scaling → PCA (≤ 40 components) → neighbor graph (15 neighbors, ≤ 30 PCs) → Leiden. The Leiden resolution was chosen from a small grid ({0.8, 1.5}) using the number of labeled types as a target cardinality hint, with the same rule applied to every arm. This uses label cardinality at evaluation time and can both inflate absolute ARI and compress differences between arms; it does not favor any one gene-selection arm.

The primary metric is mean ARI against cell_type over two Leiden seeds. Evaluation noise is of the same order as the reported effect: within-arm absolute seed-to-seed ARI differences have median ≈ 0.011 (mean ≈ 0.018), and absolute seed gaps in the shipped-path ΔARI have median ≈ 0.018, compared with a median effect of +0.016. Per-dataset deltas should therefore be read as noisy, and inference should rest on panel-level signed-rank evidence. Bootstrap percentile intervals for the median use 10,000 dataset resamples. The secondary multi-metric suite and the multiple-testing policy are described in Supplementary Note S1.

### 3.6 Ablations

Arm decomposition and *k*-move strata, cell-number baselines, the Python residual/BH FDR length baseline with its frozen-ranking control, leave-one-out threshold re-calibration, fixed-*k* rank fusion, and the external triku comparison were each run under the downstream protocol of Section 3.5. triku-specific settings are given in Section 2.8. Scripts and result tables are listed in Section 5.

### 3.7 Software

scFair v0.7.0: https://github.com/leelieber2025/scFair; https://scfair.readthedocs.io/; https://pypi.org/project/scfair/.

### 3.8 Computational cost

auto_n rebuilds an intermediate HVG-2000 → PCA → neighbors → Leiden → density pipeline once per seed (three seeds by default), so wall time is dominated by that feature pass rather than by the final ranking. On a single-core probe (OMP/MKL/OPENBLAS_NUM_THREADS=1), representative timings were: Villani (1,123 cells) 6 s for the structure pass versus 2 s for HVG@2000; Zhengmix4eq (3,994 cells) 32 s; Baron pancreas (8,569 cells) 75 s. Full automatic selection (structure plus final ranking) is of the same order. Cost therefore scales like a few standard scanpy neighbor and Leiden passes. Million-cell atlases are outside the present timing probe; for very large maps users may prefer a fixed n_top_genes, balance_method=“none”, or a *k* estimated offline on a subsample. Append itself is negligible, being set arithmetic on an existing ranked list.

## 4. Discussion

HVG list length is still set by habit. Under a frozen seurat_v3 ranking, the habitual value of 2,000 is ARI-optimal on only 1 of 18 panel datasets and leaves a mean oracle gap of +0.033. That observation stands on its own, independently of any particular rule for closing the gap, and it is the reason we treat cardinality as a design axis rather than a hyperparameter: a quantity that is wrong this often, and that every pipeline must nevertheless specify, deserves a default derived from the data. scFair supplies one that leaves the ranking untouched, and on the calibrated panel it recovers roughly half of the oracle headroom (median ΔARI = +0.016; P = 0.0077). Benchmarked end-to-end against triku at author defaults, it performs better on 15/18 datasets (median +0.024; P = 0.004), while triku itself does not beat classical HVG@2000 (median −0.018)—although the two methods act on different levers, so this compares default paths rather than a single mechanism.

The decomposition is what defines the contribution. The aggregate effect is a mixture of two arms acting on disjoint subsets, and the arms are not equally well supported. Where geometry does not move k—the majority—the path reduces to seurat_v3 top-2200, and the benefit is that a slightly longer list is adopted automatically without requiring manual tuning. Where geometry does move k, sizing carries the effect and the tail is inert. Stating this plainly matters, because it defines what a user obtains from the pipeline and where the method could be replaced by something simpler.

That raises the obvious alternative: why not simply raise the global default to 2,200? Across the full panel, HVG@2200 is not significantly better than HVG@2000 (median +0.0045; P = 0.17), and on movers it is close to neutral (+0.003), whereas the shipped path exceeds it by a median of +0.07 on that stratum. The separation is sharpest on the four short-k movers (median +0.029; range −0.003 to +0.063), where a fixed 2,200 cannot produce the 500-gene list the geometry requests. This is the mechanistic core of the method and also its narrowest evidence: two pancreas objects dominate, one non-pancreas case is weakly positive, one is a spurious short list, and the independent holdout confirms the stayer branch but not this one. The short-list advantage should thus be read as hypothesis-generating on the present panel.

Three controls clarify the interpretation of what auto_n contributes. Cell-number-only tables do not beat the classical default and systematically lengthen lists on large but compact matrices—precisely the regime where geometry shortens them—consistent with arrangement rather than count being the informative variable. The Python residual-variance FDR length rule is worse than the fixed default when its own gene content is used, but flat when only its chosen length is imposed on a frozen seurat_v3 ranking; thus, automatic length per se is not inherently beneficial, and the apparent harm stemmed from ranking content. Fixed-k rank fusion matches the shipped path’s median with a substantially heavier left tail, which is why we retain a single standard ranking rather than an ensemble. Together, these controls place our claim precisely: the value lies in how k is chosen from the geometry, not merely in the fact that k is chosen automatically. The external comparison adds one qualification—triku wins on Baron pancreas, one of the two datasets carrying the short-k advantage, demonstrating that content-based selection can outperform geometric resizing exactly where resizing appears strongest. Ranking content and list cardinality remain separable levers, and score-oriented methods remain the right tool when the ranking itself is the failure mode.

Framed more generally, the contribution is an input rather than a score. Automatic list length has previously been derived either from a statistical threshold on a per-gene statistic or from metadata such as cell number or tissue; scFair instead reads the arrangement of the cells themselves in an intermediate embedding of the matrix at hand, and lets valley depth, core count, and multi-seed stability decide how many features that arrangement requires. Letting observed structure fix a pipeline setting is well established elsewhere in the single-cell workflow—subsampling and silhouette criteria for clustering resolution in chooseR ^28^, multi-resolution cluster trees in clustree ^29^, and significance testing of cluster splits in sc-SHC ^30^ and CHOIR ^31^—but it has been applied downstream of feature selection (to the number of clusters) rather than upstream (to the number of features). Our evidence that geometry is the operative signal is exclusionary rather than direct: cell count (Section 2.10), automatic length in itself (Section 2.11), and ranking content at fixed k (Section 2.7) are each ruled out, but we did not ablate the geometric features themselves, and permuting or removing individual statistics is the obvious next test. Because this input is orthogonal to the ranking, the same features could in principle sit on top of any per-gene score, and could inform other cardinality-like settings such as the number of principal components or neighbors.

### Limitations

(i) *Calibration overlap.* Thresholds were developed with feedback from datasets overlapping panel-18. Leave-one-out re-selection never departs from the shipped constants, which is compatible with stability and with panel-fitted constants alike; only half the panel exercises the tuned branch, and the confidence guards were not re-tuned. Test-5 (*n* = 5) is under-powered and validates only the stayer branch. (ii) *Post hoc strata.* The *k*-move split was defined after the fact (8 versus 10 datasets) and the append-by-stratum tests are borderline (two-sided *P* ≈ 0.08). (iii) *Narrow short-k evidence.* Differentiation from a fixed 2,200 default rests on four post hoc datasets with limited tissue diversity, one of which (Baron pancreas) is also a dataset on which triku outperforms the shipped path. (iv) *Baselines.* The triku comparison is end-to-end at package defaults, without hyperparameter search and without separating length from content; it answers which default path wins on panel-18, not whether triku’s gene list would beat scFair at matched *n*. The FDR control is a Python residual-variance plus Benjamini–Hochberg length rule used to isolate cardinality from ranking content; it is not a package-faithful scran::getTopHVGs benchmark. (*v*) *Evaluation noise.* Primary ARI is averaged over two Leiden seeds. Within-arm seed-to-seed |ΔARI| is of the same order as the reported median effect, so per-dataset deltas are noisy and inference rests on panel-level signed-rank tests rather than single-dataset point estimates. (vi) *Label dependence.* Curated cell_type labels are themselves analysis products, resolution selection uses their cardinality, and n_types gates one guard when labels are supplied. (vii) *Scope.* Primary claims are limited to Leiden–label agreement and the secondary multi-metric suite; differential expression, marker recovery, and million-cell atlas scaling are outside this study’s endpoints. User-directed marker inclusion is offered as a manual rescue for unresolved populations and is not itself a benchmarked method.

Practical recommendations follow directly. Use the automatic path for exploratory analysis; fix an explicit integer n for published figures; set balance_method=“none” when an exact list length is required; and do not substitute a global default of 2,200 without checking whether the dataset falls in a short-list regime.

## 5. Data and code availability

Data availability. All count matrices analyzed in this study were obtained from public repositories; no private laboratory datasets were used. The primary evaluation panel (panel-18; n = 18) comprises three groups.

### (i) FACS/sort references

Zhengmix4eq, Zhengmix4uneq, and Zhengmix8eq purified-PBMC mixtures from the DuoClustering2018 Bioconductor package ^18^, derived from Zheng et al. ^17^ and redistributed in the P1 HVG benchmark archive (Zenodo https://doi.org/10.5281/zenodo.12135988); a nine-population FACS PBMC panel from Zheng et al. ^17^; blood dendritic-cell and monocyte Smart-seq2 profiles (GEO: GSE94820) ^19^; and Tabula Muris FACS limb-muscle and brain myeloid/non-myeloid plates (figshare https://doi.org/10.6084/m9.figshare.5829687; GEO: GSE109774)^21^.

### (ii) Author-labeled or multi-protocol objects

human pancreatic islets (GEO: GSE84133) ^26^; mouse small-intestine epithelium (GEO: GSE92332) ^22^; multi-protocol human pancreas and lung integration datasets from the scIB atlas-level benchmark (figshare https://doi.org/10.6084/m9.figshare.12420968) ^23^; glioblastoma Smart-seq2 profiles (GEO: GSE84465) ^24^; and the unperturbed three-cell-line mixture underlying CraftedExperiment (GEO: GSE136148) ^7^.

### (iii) Commercial and protein-annotated references

10x Genomics 10k PBMC 34 v3 gene expression (https://www.10xgenomics.com/datasets/10-k-pbm-cs-from-a-healthy-donor-v-3-chemistry-3-standard-3-0-0) with eight-type author labels; 10x Genomics 10k PBMC CITE-seq with TotalSeq-B surface protein (https://www.10xgenomics.com/datasets/10-k-pbm-cs-from-a-healthy-donor-gene-expression-and-cell-surface-protein-3-standard-3-0-0) with ADT-mapped type labels; Seurat v4 multimodal PBMC CITE-seq (GEO: GSE164378; https://atlas.fredhutch.org/nygc/multimodal-pbmc/) ^20^; mouse spleen and lymph-node CITE-seq SLN-208 as released with totalVI ^25^; and a CELLxGENE Discover curated 10x v3 PBMC object (https://cellxgene.cziscience.com/). The independent holdout (Test-5; five organs) is drawn from Tabula Sapiens ^16^ via CELLxGENE Discover (collection https://cellxgene.cziscience.com/collections/e5f58829-1a66-40b5-a624-9046778e74f5), restricted to 10x Chromium 34 v3 chemistry and consortium manually annotated Cell Ontology labels.

## Code availability

scFair v0.7.0 is available from GitHub (https://github.com/leelieber2025/scFair), PyPI (https://pypi.org/project/scfair/), and Read the Docs (https://scfair.readthedocs.io/). Analyses reported here correspond to scFair v0.7.0, git commit 7db77dba81e9816d165713bc8befeabcadeba0fb. Scripts that regenerate the reported tables and figures, together with frozen summary result tables, are included in that public repository. A Zenodo archive of frozen result tables and version-tagged releases is available at https://doi.org/10.5281/zenodo.21761251 (concept DOI; resolves to all versions).

## Author contributions

Z.L. conceived the method, implemented the software, designed and performed the analyses, and wrote the manuscript. A.W.J. edited the manuscript. SX.L. implemented the software.

## Acknowledgements

The author thanks the generators of the public datasets used for validation.

## References

1. Brennecke, P., et al. Accounting for technical noise in single-cell RNA-seq experiments. Nat Methods 10, 1093–1095 (2013).

2. Satija, R., Farrell, J. A., Gennert, D., Schier, A. F. & Regev, A. Spatial reconstruction of single-cell gene expression data. Nat Biotechnol 33, 495–502 (2015).

3. Stuart, T., et al. Comprehensive Integration of Single-Cell Data. Cell 177, 1888–1902.e21 (2019).

4. Wolf, F. A., Angerer, P. & Theis, F. J. SCANPY: large-scale single-cell gene expression data analysis. Genome Biology 19, 15 (2018).

5. Zhao, R., et al. A systematic evaluation of highly variable gene selection methods for single-cell RNA-sequencing. Genome Biol 26, 424 (2025).

6. Yip, S. H., Sham, P. C. & Wang, J. Evaluation of tools for highly variable gene discovery from single-cell RNA-seq data. Brief Bioinform 20, 1583–1589 (2019).

7. Liu, S., Corcoran, D. L., Garcia-Recio, S., Marron, J. S. & Perou, C. M. Crafted experiments to evaluate feature selection methods for single-cell RNA-seq data. NAR Genom Bioinform 7, lqaf023 (2025).

8. Dollinger, E. P., Silkwood, K., Atwood, S., Nie, Q. & Lander, A. D. Statistically principled feature selection for single cell transcriptomics. BMC Bioinformatics 26, 238 (2025).

9. Luecken, M. D. & Theis, F. J. Current best practices in single-cell RNA-seq analysis: a tutorial. Mol Syst Biol 15, e8746 (2019).

10. Hafemeister, C. & Satija, R. Normalization and variance stabilization of single-cell RNA-seq data using regularized negative binomial regression. Genome Biol 20, 296 (2019).

11. Lun, A. T. L., McCarthy, D. J. & Marioni, J. C. A step-by-step workflow for low-level analysis of single-cell RNA-seq data with Bioconductor. F1000Res 5, 2122 (2016).

12. Andrews, T. S. & Hemberg, M. M3Drop: dropout-based feature selection for scRNASeq. Bioinformatics 35, 2865–2867 (2019).

13. Townes, F. W., Hicks, S. C., Aryee, M. J. & Irizarry, R. A. Feature selection and dimension reduction for single-cell RNA-Seq based on a multinomial model. Genome Biol 20, 295 (2019).

14. M Ascensión, A., Ibáñez-Solé, O., Inza, I., Izeta, A. & Araúzo-Bravo, M. J. Triku: a feature selection method based on nearest neighbors for single-cell data. Gigascience 11, giac017 (2022).

15. Traag, V. A., Waltman, L. & van Eck, N. J. From Louvain to Leiden: guaranteeing well-connected communities. Sci Rep 9, 5233 (2019).

16. Tabula Sapiens Consortium* et al. The Tabula Sapiens: A multiple-organ, single-cell transcriptomic atlas of humans. Science 376, eabl4896 (2022).

17. Zheng, G. X. Y., et al. Massively parallel digital transcriptional profiling of single cells. Nat Commun 8, 14049 (2017).

18. Duò, A., Robinson, M. D. & Soneson, C. A systematic performance evaluation of clustering methods for single-cell RNA-seq data. F1000Res 7, 1141 (2018).

19. Villani, A.-C., et al. Single-cell RNA-seq reveals new types of human blood dendritic cells, monocytes, and progenitors. Science 356, eaah4573 (2017).

20. Hao, Y., et al. Integrated analysis of multimodal single-cell data. Cell 184, 3573–3587.e29 (2021).

21. Tabula Muris Consortium et al. Single-cell transcriptomics of 20 mouse organs creates a Tabula Muris. Nature 562, 367–372 (2018).

22. Haber, A. L., et al. A single-cell survey of the small intestinal epithelium. Nature 551, 333–339 (2017).

23. Luecken, M. D., et al. Benchmarking atlas-level data integration in single-cell genomics. Nat Methods 19, 41–50 (2022).

24. Darmanis, S., et al. Single-Cell RNA-Seq Analysis of Infiltrating Neoplastic Cells at the Migrating Front of Human Glioblastoma. Cell Rep 21, 1399–1410 (2017).

25. Gayoso, A., et al. Joint probabilistic modeling of single-cell multi-omic data with totalVI. Nat Methods 18, 272–282 (2021).

26. Baron, M., et al. A Single-Cell Transcriptomic Map of the Human and Mouse Pancreas Reveals Inter- and Intra-cell Population Structure. Cell Syst 3, 346–360.e4 (2016).

27. Wang, F. & Rudin, C. Falling Rule Lists. in Proceedings of the Eighteenth International Conference on Artificial Intelligence and Statistics 1013–1022 (PMLR, 2015).

28. Patterson-Cross, R. B., Levine, A. J. & Menon, V. Selecting single cell clustering parameter values using subsampling-based robustness metrics. BMC Bioinformatics 22, 39 (2021).

29. Zappia, L. & Oshlack, A. Clustering trees: a visualization for evaluating clusterings at multiple resolutions. Gigascience 7, giy083 (2018).

30. Grabski, I. N., Street, K. & Irizarry, R. A. Significance analysis for clustering with single-cell RNA-sequencing data. Nat Methods 20, 1196–1202 (2023).

31. Sant, C., Mucke, L. & Corces, M. R. CHOIR improves significance-based detection of cell types and states from single-cell data. Nat Genet 57, 1309–1319 (2025).

