## Supplemental for "scFair: Geometry-Aware Gene Budgets and Same-Rank Extension for Highly Variable Gene Selection"

### **Supplementary Information**

#### Supplementary figures

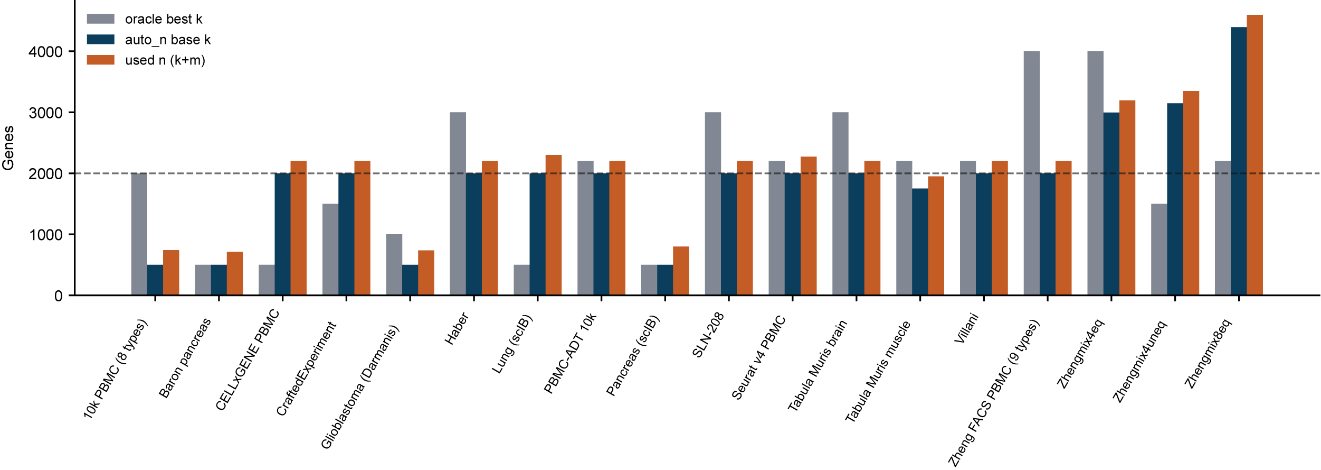

**Supplementary Figure S1.** Full gene budgets on panel-18 (oracle best *k*, auto_n base *k*, and used *n* = *k* + *m*).

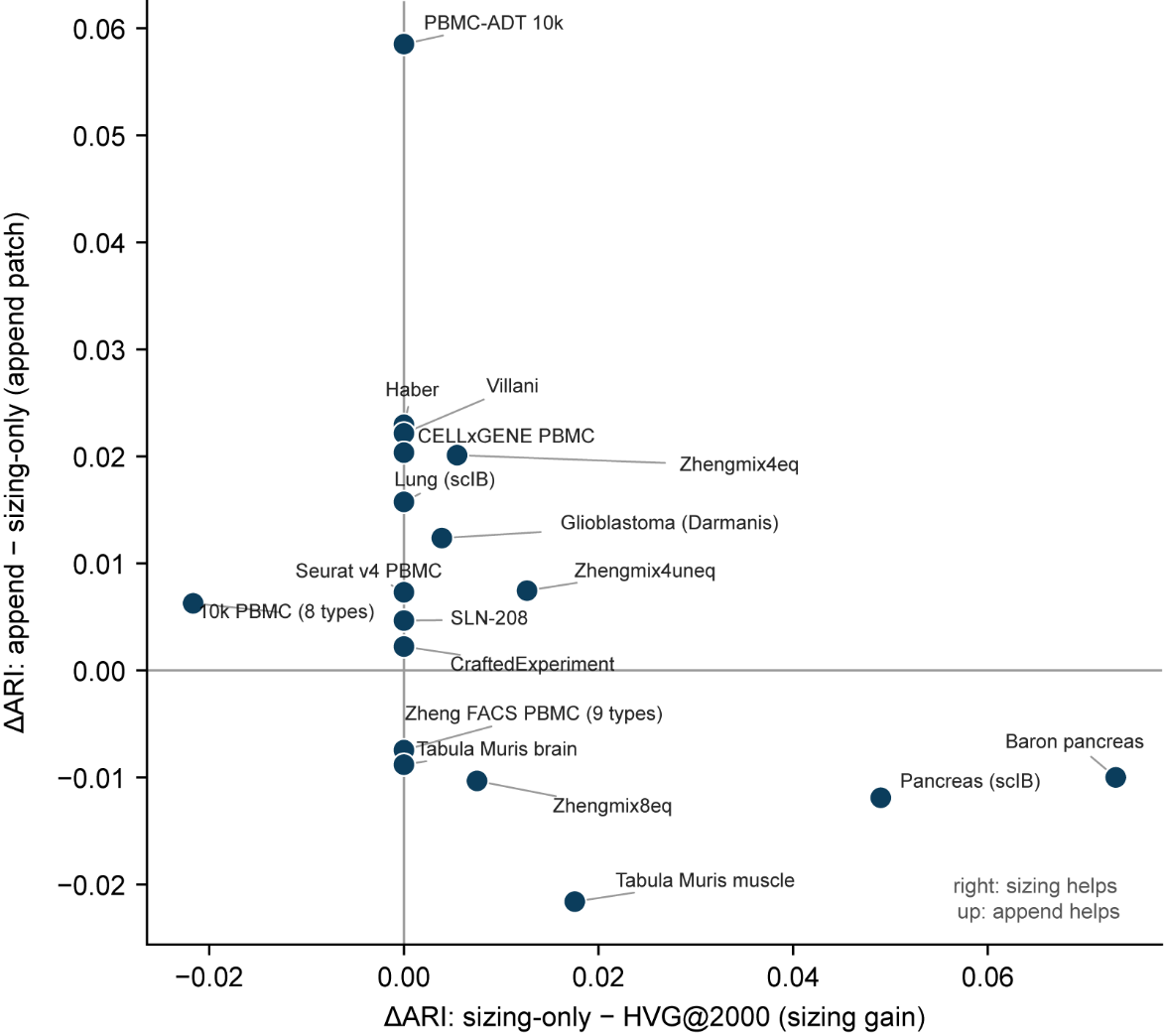

**Supplementary Figure S2.** Per-dataset sizing gain (sizing-only − HVG@2000) versus append patch (append − sizing-only) on panel-18.

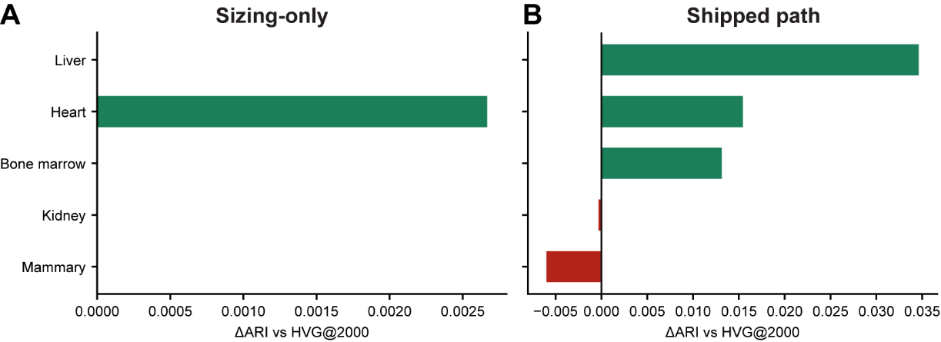

**Supplementary Figure S3.** Tabula Sapiens organ holdout (Test-5): sizing-only and the shipped path versus HVG@2000, per organ.

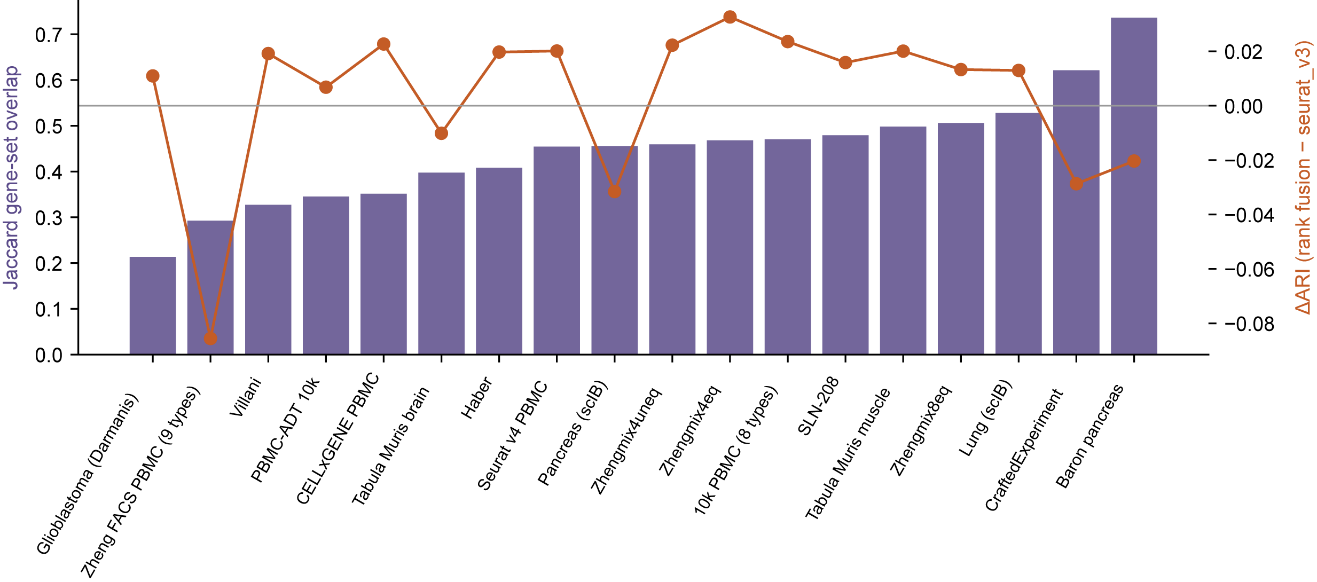

**Supplementary Figure S4.** Rank fusion changes gene-set membership (Jaccard) without robust ARI gains at fixed *k* (panel-18).

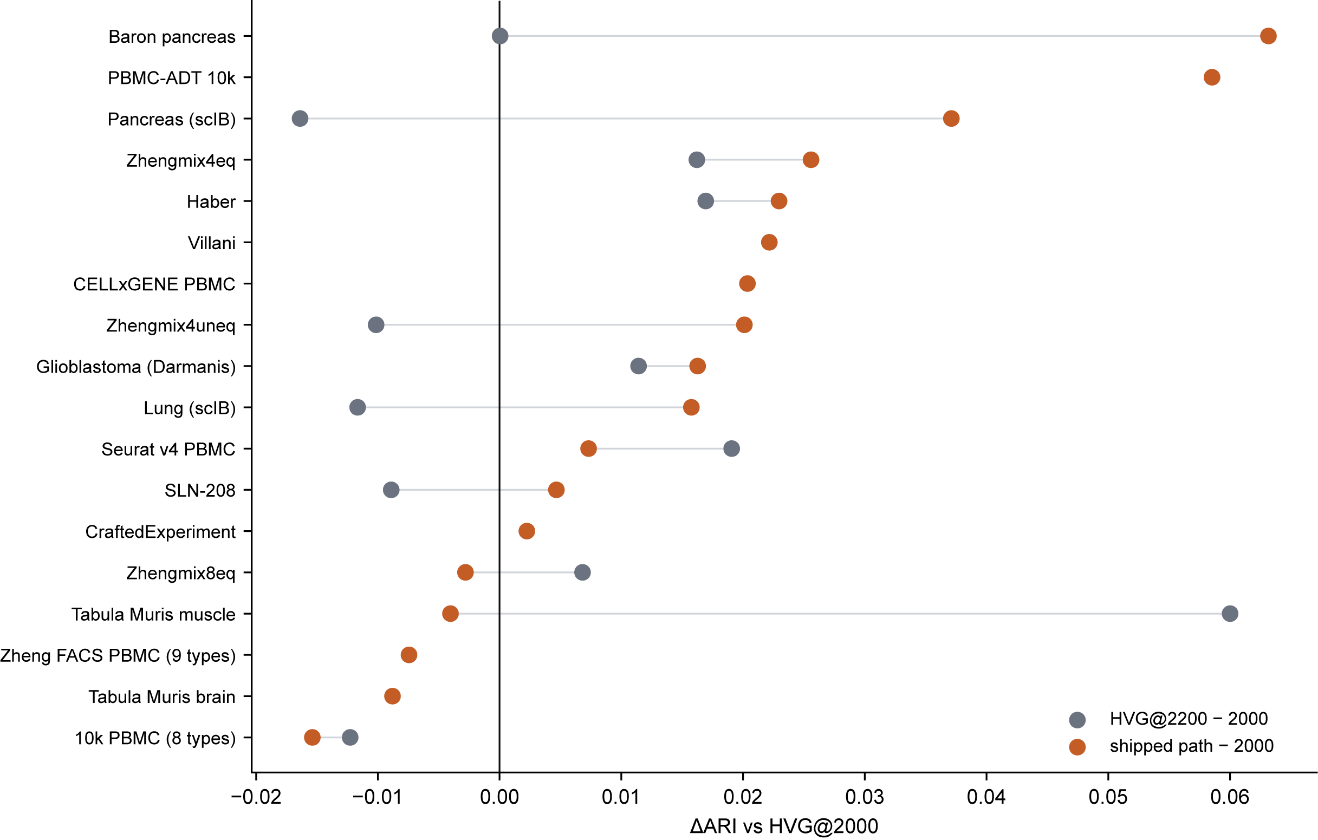

**Supplementary Figure S5.** Shipped-path ΔARI versus the fixed HVG@2200 control, each relative to HVG@2000 (panel-18).

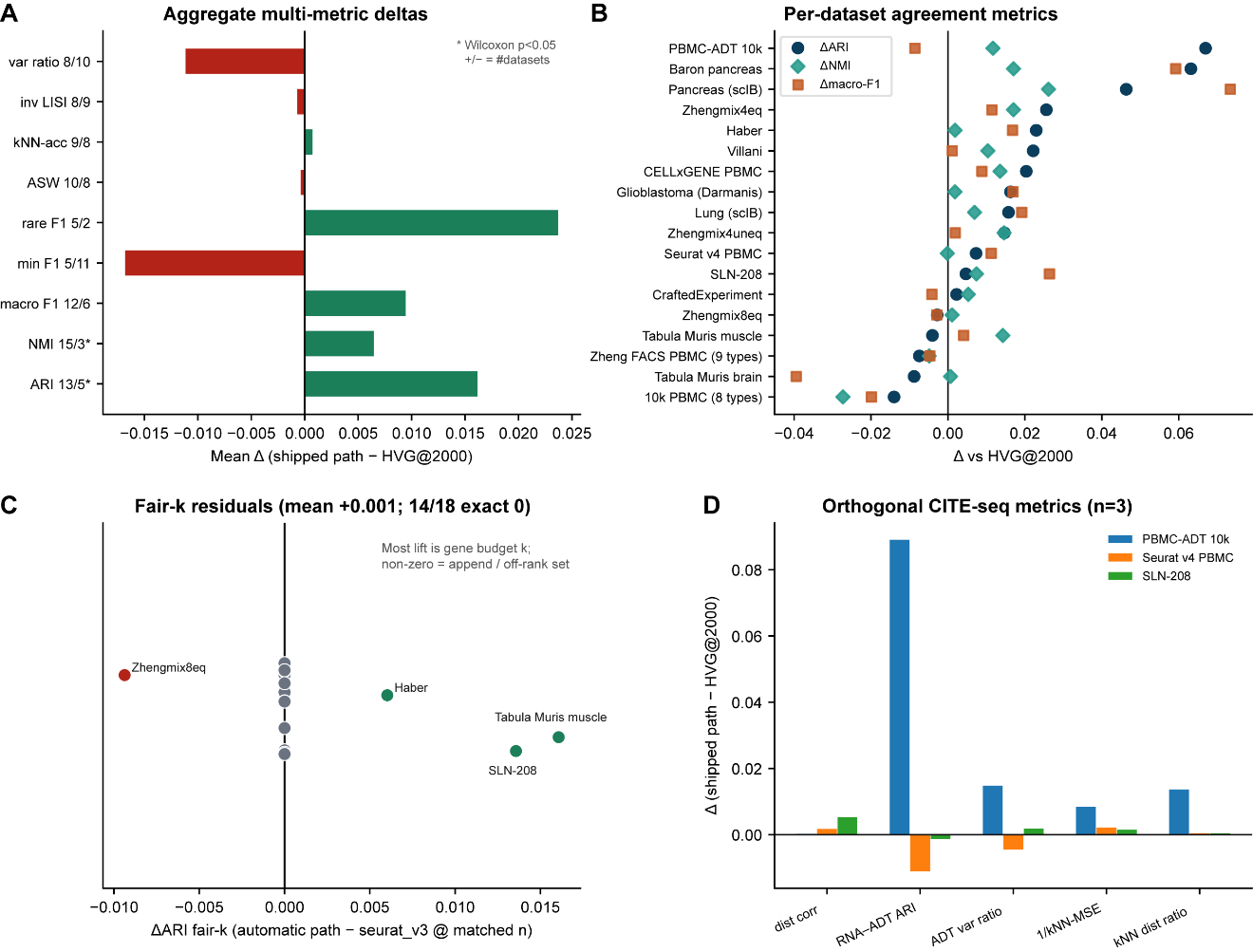

**Supplementary Figure S6.** Multi-metric evaluation on panel-18. (**A**) Mean shipped − HVG@2000 deltas across metrics (asterisk, Wilcoxon *P* < 0.05; positive/negative dataset counts). (**B**) Per-dataset ΔARI, ΔNMI, and Δmacro-F1. (**C**) Fair-*k* residuals (automatic path versus seurat_v3 at matched *n*): 14/18 are exactly zero, which reflects gene-set identity wherever the selected set is the seurat_v3 top-*n* list (see Note S1) rather than independent evidence about *k*. (**D**) Orthogonal CITE-seq metrics on three protein-paired objects.

#### Supplementary Note S1. Multi-metric suite

Primary claims use Leiden ARI against curated cell_type labels. To address metric gaps raised by feature-selection benchmarks — partition purity, minority-type recovery, and ranking-orthogonal protein evidence — we re-ran the shipped protocol against HVG@2000 on panel-18 with two seeds, adding a fair-*k* control (seurat_v3 at the selected gene count).

Agreement metrics track the primary result: median ΔARI = +0.016 (mean +0.0154; 13/5; *P* = 0.0077) and mean ΔNMI = +0.0065 (15/3; *P* = 0.0066). Embedding-geometry proxies are flat (mean ΔASW ≈ −0.0004, *P* = 0.58; Δ*k*-NN accuracy ≈ +0.0008, *P* = 0.71; Δinverse-LISI ≈ −0.0008, *P* = 0.93). Type-level F1 is mixed. The fair-*k* control has mean ΔARI = +0.0015 with 14/18 exact zeros; those zeros are expected whenever the selected set is exactly the seurat_v3 top-*n* list, which holds for most but not all *m* = 200 stayers at *n* = 2,200. Fair-*k* identity is therefore a statement about gene-set equality, not independent evidence that only *k* matters beyond the ranking. The four non-zero fair-*k* residuals — Haber, SLN-208, Zhengmix8eq, and Tabula Muris muscle — are not explained by density-raised tails (both density-raised stayers are exact zero) or by mover status alone (6/8 movers are exact zero); the common cause is an exact tie in the seurat_v3 score at that dataset's selected-list cutoff, where scFair's rank construction and scanpy's native tie-break diverge (Methods 3.4).

*Multiple testing.* Only the panel-18 ΔARI contrast against HVG@2000 is treated as primary. Applying Benjamini–Hochberg control at *q* = 0.05 across the nine label-space mean deltas (ARI, NMI, ASW, *k*-NN accuracy, variance ratio, inverse LISI, rare-F1, macro-F1, minimum-F1) retains ARI and NMI; the remaining geometry and F1 contrasts are not discoveries. CITE-seq tests (*n* = 3) are descriptive only.

*Orthogonal protein readouts.* On three CITE-seq objects with paired protein measurements (PBMC-ADT 10k, Seurat v4 20k PBMC, SLN-208 mouse), the shipped path versus HVG@2000 gives small non-negative mean deltas for protein distance correlation (+0.0025; 3/3), inverse *k*-NN MSE (+0.0041; 3/3), and *k*-NN distance ratio (+0.0049; 3/3); RNA–ADT Leiden ARI is mixed. These readouts do not replace ARI as the primary endpoint but bound over-interpretation of geometry-only improvements.

#### Supplementary Table S1. The auto_n decision list (shipped structure rule)

Table S1a gives the ordered, first-match-wins decision list *R* = [(*c*₁,*k*₁), …, (*c*₉,*k*₉)] that assigns the raw base size *k*₀ from the seed-aggregated structure statistics (*v* = valley median, *f* = fraction shallow, *n*_d = number of density cores, μ_s = mean stability, μ_s,min = minimum stability, *r* = *n*_leiden / *n*_d). Rows are evaluated in order and the first matching condition determines *k*₀. Equation (1) in Methods 3.3 gives the closed form for row 1; all other rows assign one of four fixed values. *k*_min = 500, *k*_max = 5,000, and *k*₀ is further capped at the number of genes assayed. Regime names describe the geometric assignment rather than software identifiers.

| # | Condition | *k*₀ | Regime |
| --- | --- | --- | --- |
| 1 | *f* ≥ 0.85 and *n*_d ≤ 4.5 | Eq. (1) | Long (shallow valleys, few cores) |
| 2 | *n*_obs ≥ 10,000 and *n*_d ∈ [12,20] and *v* ≥ 0.78 and *r* ≤ 1.12 | 2000 | Fine-structure / atlas band |
| 3 | *v* ≥ 0.80 and *n*_d ≥ 6 | 500 | Short-hard (deep valleys, ≥6 cores) |
| 4 | *v* ≥ 0.70 and *n*_d ≥ 12 | 500 | Short-hard (moderate valleys, ≥12 cores) |
| 5 | *v* ≥ 0.65 and 6 ≤ *n*_d < 12 | 1500 (2000 if μ_s,min < 0.1 and μ_s < 0.55) | Mid / Mid raised when unstable |
| 6 | μ_s,min < 0.25 and *n*_d ≤ 5, and (*f* < 0.85 or μ_s,min < 0.2) | 500 | Short-unstable (coarse geometry) |
| 7 | *v* ≥ 0.65 | 1000 | Soft mid-short (1,000; valley depth) |
| 8 | μ_s < 0.45 and *v* ≥ 0.5 | 1000 | Soft mid-short (1,000; low stability) |
| 9 | otherwise | 2000 | Classical default |

**Table S1b** **lists, in order, the confidence guards applied to the Table S1a match**. Final *k* = clip(*k*₀ after guards, *k*_min, *k*_max, *n*_genes).

| # | Guard | Trigger | Effect |
| --- | --- | --- | --- |
| 1 | Soft ladder buffer | raw *k*₀ ∈ {500, 1000, 1500} | one rung up: 500→1000, 1000→1500, 1500→2000 |
| 2 | True short-list exception | short-hard regime, *n*_d ≥ 10, *n*_types ≥ 5 (labels only) | buffer skipped; *k* stays 500 |
| 3 | Spurious short-list floor | short-hard regime, *n*_obs ≥ 10,000, low density confidence, *n*_d ≤ 8 | *k* floored to 2000 |
| 4 | Residual anti–short-list | *k* ≤ 500 with untrusted density (low confidence, high depth sensitivity, or large *n* with few cores) | *k* floored to 2000 |
| 5 | Low-confidence floor | *k* < 2000 and low density confidence | *k* floored to 2000 (true short-list *k* = 500 exempt) |
| 6 | Fine-mode floor | fine HVG mode is requested and *k* < 2000 on short or soft regimes | *k* floored to 2000 |

#### Supplementary Table S2. Design axes for automatic or thresholded HVG list length

| Approach | What sets the list length | Uses embedding geometry of the current matrix | Ranking held fixed | Ecosystem |
| --- | --- | --- | --- | --- |
| scanpy n_top_genes ^4^ | User-supplied integer (habitually 2,000) | No | Yes | Python |
| scran getTopHVGs ^11^ | FDR or residual-variance threshold on the per-gene statistic | No | No (length is tied to the scran statistic) | R/Bioconductor |
| M3Drop ^12^ | Dropout-model FDR threshold | No | No | R |
| Deviance-based selection ^13^ | User-supplied integer on a different statistic | No | No | R |
| triku ^14^ | Neighborhood-expression score threshold | Uses a *k*-NN graph, not density regime | No | Python |
| Cell-count rules (Section 2.10) | Step table or √*n*_obs on cell number | No | Yes | — |
| scFair auto_n (this work) | Multi-seed density/stability regime, with floors to 2,000 under weak evidence | Yes | Yes | Python / scanpy |

Conceptual comparison of how list length is chosen; not a multi-package benchmark. End-to-end quantitative comparison against an external selector is limited to triku (Section 2.8; triku 2.2.0, author defaults, no tuning: triku − HVG@2000 median ΔARI = −0.018, 13/18 negative; shipped − triku median +0.024, 15/18, P = 0.004). Cell-count and FDR-length controls are reported in Sections 2.10 and 2.11; the FDR-length control is the Python residual/BH rule of Section 2.11, not Bioconductor getTopHVGs. Entries for scran, M3Drop, and deviance-based selection locate design differences only.
